# Phosphorylation-inducing molecules for regulating dynamic cellular processes

**DOI:** 10.1101/2025.05.10.652895

**Authors:** Rajaiah Pergu, Sreekanth Vedagopuram, Praveen Kokkonda, Sophia Lai, Praveen K. Tiwari, Sameek Singh, Junya Kawai, Santanu Singha, Wenzhi Tian, Vikram Thimaradka, Kien Tran, Amit Choudhary

**Affiliations:** Chemical Biology and Therapeutics Science, Broad Institute of MIT and Harvard, Cambridge, MA 02142, USA; Department of Medicine, Harvard Medical School, Boston, MA 02115, USA; Divisons of Renal Medicine and Engineering, Brigham and Women’s Hospital, Boston, MA 02115, USA; Department of Chemistry and Chemical Biology, Harvard University, 12 Oxford Street, Cambridge, MA 02138, USA

## Abstract

Dynamic cellular processes often employ protein phosphorylation for rapid information transfer within and between cells. Phosphorylation-inducing chimeric small molecules (PHICS) have been developed for targeted protein phosphorylation by on-demand inducing a kinase-protein pairing. However, widespread application of PHICS has been limited as previously reported PHICS that recruited AMP-activated protein kinase (AMPK) required serum starvation and target protein overexpression, recruited only a few of the potential AMPK complexes, and exhibited poor dose- and temporal control. Herein, we report an AMPK PHICS platform that operates under physiological conditions (i.e., no serum starvation or target overexpression), recruits multiple AMPK complexes, and can induce target protein phosphorylation with dose and temporal control. We demonstrated the utility of this platform for controlling phosphorylations that underlie two dynamic cellular processes, namely oncogenic signaling and phase separation. An AMPK PHICS directed against Bruton’s Tyrosine Kinase (BTK), which is a driver of several B cell malignancies, was effective at inducing the death of drug-resistant cancer cells. Here, PHICS induced inhibitory phosphorylations on BTK and attenuated its oncogenic pathway. Phosphorylation of Liprin-α3 induces phase separation in neurons and is critical for neurotransmitter release. Using AMPK PHICS that phosphorylated Liprin-α3, we induced Liprin-α3 phase separation with dose and temporal control. We envision this PHICS platform to find utility in inducing and controlling protein phosphorylations for basic research and biomedicine.

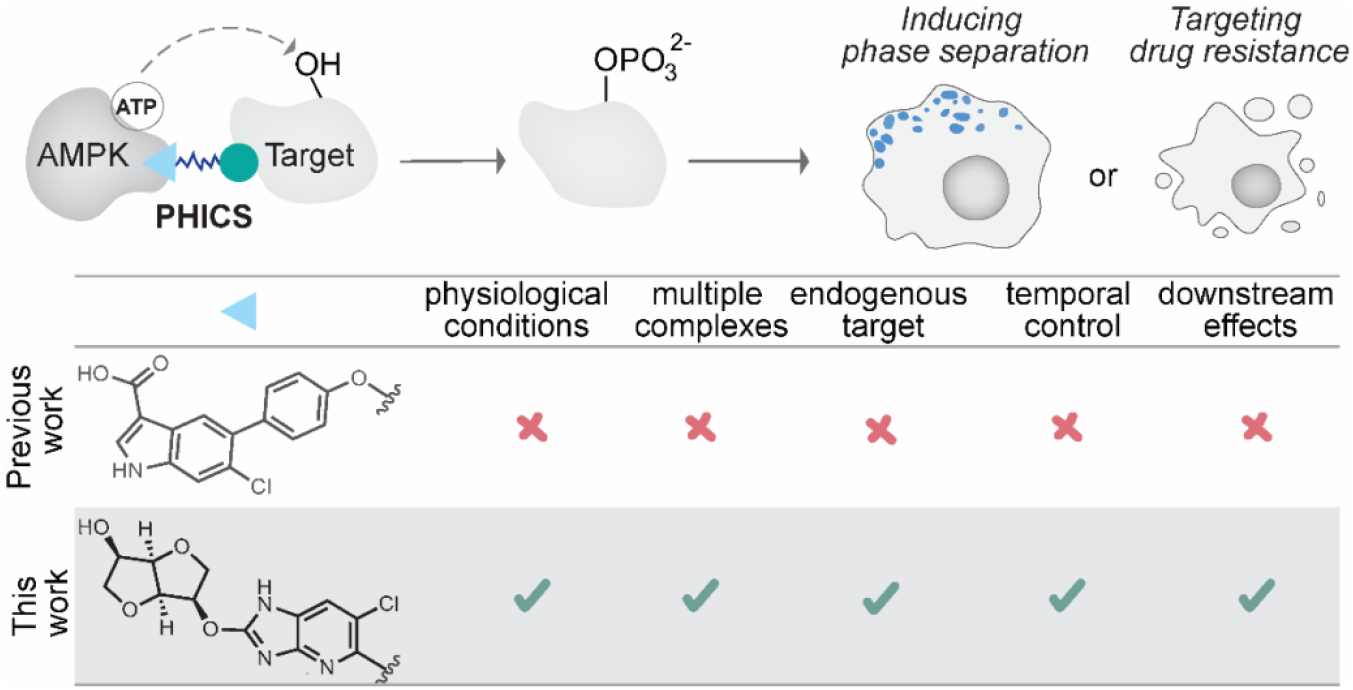

## Introduction

Nature employs posttranslational modifications (PTMs) to regulate highly dynamic processes using specific enzymes that either “write” or “erase” PTMs on a target protein.^1, 2^ Leveraging the ability of small molecules that afford dose and temporal control of enzymic activity,^3^ several groups have reported synthetic chimeric molecules for writing or erasing PTMs. These chimeric molecules,^4, 5^ formed by connecting binders of a target and an endogenous PTM-writing or -erasing enzyme, induce proximity between the enzyme and target, resulting in PTM induction or removal even when the target protein is not the enzyme’s natural substrate. For example, proteolysis-targeting chimeras (PROTACs)^6^ have been developed to induce proximity between the target protein and E3 ubiquitin ligase, causing ubiquitination and subsequent protein degradation. Chimeras have also been reported for writing or erasing PTMs, such as deubiquitinase-targeting chimeras (DUBTACs) that remove ubiquitin,^7^ phosphorylation-targeting chimeras (PhosTACs) and phosphatase-targeting chimeras (PHORCs) that remove phosphoryl group,^8, 9^ heterobifunctional molecules that induce acetylation (AceTAGs),^10^ and bifunctional nanobody conjugates that add or remove glycosylation.^11, 12^

We recently described phosphorylation-inducing chimeric small molecules (PHICS) that allow kinases to phosphorylate target proteins, including non-natural substrates.^13-15^ These chimeras operate on the principle of chemically-induced proximity: linking binders of the kinase and target protein together can artificially induce an enzyme-substrate pairing that results in targeted protein phosphorylation. While the reported kinase binder in PHICS to recruit AMP-activated protein kinase (AMPK) performed well in proof-of-concept systems, the binder suffers from serious flaws preventing the widespread application of PHICS. For instance, previous AMPK ligands required overexpression of the target protein and serum starvation of cells, and they exhibited poor dose- and temporal control, all of which prevent PHICS applications in realistic biological scenarios. Additionally, the reported AMPK ligands recruited only β1 isoforms of AMPK, which is a complex comprising three heterotrimeric subunits, each with various isoforms—the α subunit has two isoforms, β has 2, and γ has 3, for 12 possible AMPK complexes^16^ in mammalian cells. As these complexes vary in expression across cell types, an AMPK binder that can recruit all complexes can be useful in myriad scenarios.^17^

Herein, we describe a new AMPK PHICS platform that uses a pan-AMPK binder capable of recruiting multiple AMPK complexes (**Figure 1A**) and that functions on endogenous proteins without cell starvation, allowing precise dose and temporal control of biological pathways. With this platform, we showed the first examples of AMPK-induced phosphorylation in non-native targets, including endogenous BCR-ABL, an aberrant, constitutively active fusion protein of BCR and ABL tyrosine kinase commonly expressed in CML patients.^18^ We also applied this platform to two relevant biological scenarios. Bruton Tyrosine Kinase (BTK) is a driver of the proliferation of malignant B cells in chronic lymphocytic leukemia and mantle cell lymphoma and resistance has emerged for drugs targeting BTK.^19^ We developed PHICS that induce inhibitory phosphorylations on BTK and these PHICS also impeded growth of drug-resistant cancer cell lines. The active zone in neurons, which plays a critical role in neurotransmitter release, is a highly dynamic and specialized structure controlled by phosphorylation-induced phase separation of Liprin-α3.^20^ Using AMPK PHICS targeting Liprin-α3, we induced its phase separation with dose- and temporal control. Using global phosphoproteomics and genetic methods, we identified and validated the phosphorylation site. These studies led to discovery of a Liprin-α3 variant with a two-fold higher degree of phosphorylation-mediated phase separation. Overall, these studies describe a new PHICS platform to induce serine/threonine phosphorylation with dose and temporal control that enabled induction of phase-separation triggering phosphorylation and inhibitory phosphorylation to block an oncogenic signaling pathway.

**Figure 1.**
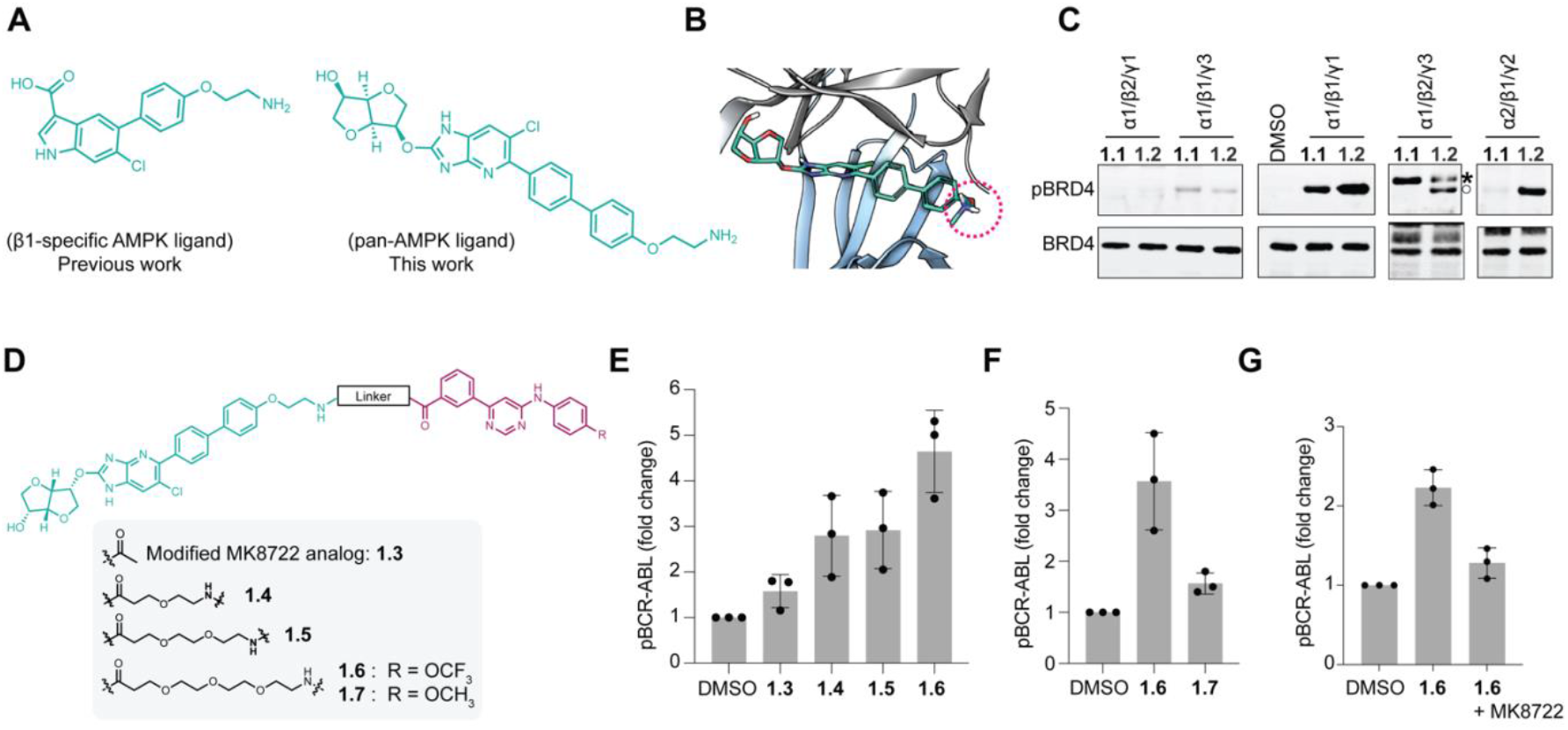
Design of AMPK PHICS. (**A**) Binders for β1-specific AMPK and pan-AMPK recruitment. (**B**) Molecular docking showing the solvent-accessible end of the modified MK8722 analog. (**C**) Treatment of BRD4-GST with **1.1** or **1.2** and five AMPK complexes revealed differences in BRD4 phosphorylation. Asterisk refers to GST-tagged AMPK subunit, and the open circle refers to the phosphorylated target protein (p-BRD4-GST). BRD4 and two of the AMPK subunits (α1/β2/γ3 and α2/β1/γ2) have GST-tag. (**D**) Chemical structures of the BCR-ABL PHICS and its analogs. (**E**) Induction of BCR-ABL phosphorylation by PHICS. (**F**) Comparison of phosphorylation-inducing activities of active and inactive PHICS analogs in K562 cells. (**G**) Competition experiment using MK8722 competition (1:1 at 5 µM) against **1.6** in K562 cells.

## Results and Discussion

### Development of AMPK PHICS platform

The efficacy of AMPK PHICS would be increased via the capacity to recruit various AMPK complexes with differing intrinsic activities. To develop a pan-AMPK PHICS, we used molecular docking of the reported pan-AMPK binder MK8722 to identify the linker attachment site as the para position of the terminal phenyl ring of MK8722 that is located on the opposite end of the isomannide ring (**Figure 1B and Figure S1A**).^21, 22^ In addition to being solvent inaccessible, the free hydroxyl on the isomannide forms hydrogen bonds to Asn48 and Ile46, which should not be perturbed to maximize binding affinity (**Figure S1B**). With the chosen attachment site, the para-appended 2-phenoxylethylamino linker emerges between the α (catalytic domain) and β subunits (**Figure S2**).

To test the ability of MK8722-based PHICS to recruit multiple isoforms of AMPK, we measured the phosphorylation induced in the target bromodomain-containing protein 4 (BRD4)^13^ in the presence of various PHICS and isoforms of AMPK. Since the AMPK complexes differ in their intrinsic kinase activities, it is not meaningful to directly compare the phosphorylation levels between the various complexes but rather to observe the relative phosphorylation induced *via* various PHICS. To thus generate a direct comparison, we synthesized two PHICS with different AMPK ligands, with **1.1** containing the β1-specific AMPK binder from our previous work and **1.2** containing our modified MK8722 (**Figure S3**). Using immunoblotting, we measured the ability of these compounds to phosphorylate recombinant truncated BRD4-GST *via* five different AMPK heterotrimers— α1/β1/γ1, α1/β2/γ1, α1/β1/γ3, α1/β2/γ3, and α2/β1/γ2 (**Figure 1C**). Comparing the relative activities of **1.1** versus **1.2**, isoforms α1/β1/γ1, α1/β2/γ1, and α1/β1/γ3 saw similar BRD4 phosphorylation levels between the two PHICS, while isoforms α1/β2/γ3, and α2/β1/γ2 only induced phosphorylation in the presence of **1.2**. Thus, we have demonstrated that this pan-specific AMPK binder can recruit more complexes to its target protein of interest, expanding the potential generalizability of the AMPK PHICS platform.

Next, to test our pan-AMPK PHICS against an endogenous protein, we targeted BCR-ABL, which is a constitutively active fusion of BCR and ABL, and driver of CML pathogenesis.^23^ To design PHICS against BCR-ABL, we used the allosteric and non-ATP competitive inhibitor GNF-2 because of its selectivity against BCR-ABL compared to other tyrosine kinases, including wild-type ABL.^24^ We generated PHICS with various linker lengths (**Figure 1D**) to identify the analog that induced the highest phosphorylation level. Among the PHICS tested, **1.6** generated the highest level of endogenous BCR-ABL phosphorylation in K562 cells (**Figure 1E, S4A**). We generated a control compound **1.7** that has reduced affinity^25^ for BCR-ABL and this control compound induced less phosphorylation (**Figure 1F, S4B**). To demonstrate whether MK8722 could compete with **1.6** and reduce phosphorylation levels, we co-treated **1.6** with MK8722 and observed reduced BCR-ABL phosphorylation (**Figure 1G, S4C**). We also observed dose-dependent phosphorylation induction of endogenous BCR-ABL phosphorylation in K562 cells by **1.6** (**Figure S4D**). Importantly, phosphorylation of BCR-ABL was observed in as little as 0.5–1 µM **1.6** without needing to overexpress either BCR-ABL or AMPK. Overall, these results suggest that PHICS can induce phosphorylation on the endogenous target protein.

To determine if the new PHICS platform can induce phosphorylation without serum starvation of cells, we used BTK as the target protein whose phosphorylation (pS180) by previous PHICS required serum starvation (**Figure S5A**).^13^ We synthesized several BTK PHICS containing various linker lengths and chemotypes (**Figure 2E**) and tested them in HEK293T cells transiently expressing BTK. From the compounds tested, the newly generated PHICS **2.3, 2.4**, and **2.5** yielded a higher phosphorylation (pS180) level as compared to previously generated **2.1** and **2.2**, though interestingly, PHICS did not require the HEK293T cells to be serum starved to induce phosphorylation (**Figure 2B, S5B, C**). In addition, the BTK PHICS generated herein induced phosphorylation on BTK (pS180) at lower concentrations than had been required in the previous work (**Figure S5B**). This might be due to the ability of a pan-AMPK binder to recruit many different AMPK complexes, making it statistically more likely to induce a ternary complex between AMPK and the target protein BTK. We note that individual components of BTK PHICS (AMPK or BTK binder) did not induce S180 phosphorylation (**Figure S5C**). Overall, these studies suggest that the AMPK PHICS platform can induce phosphorylation of target proteins without serum starvation.

**Figure 2.**
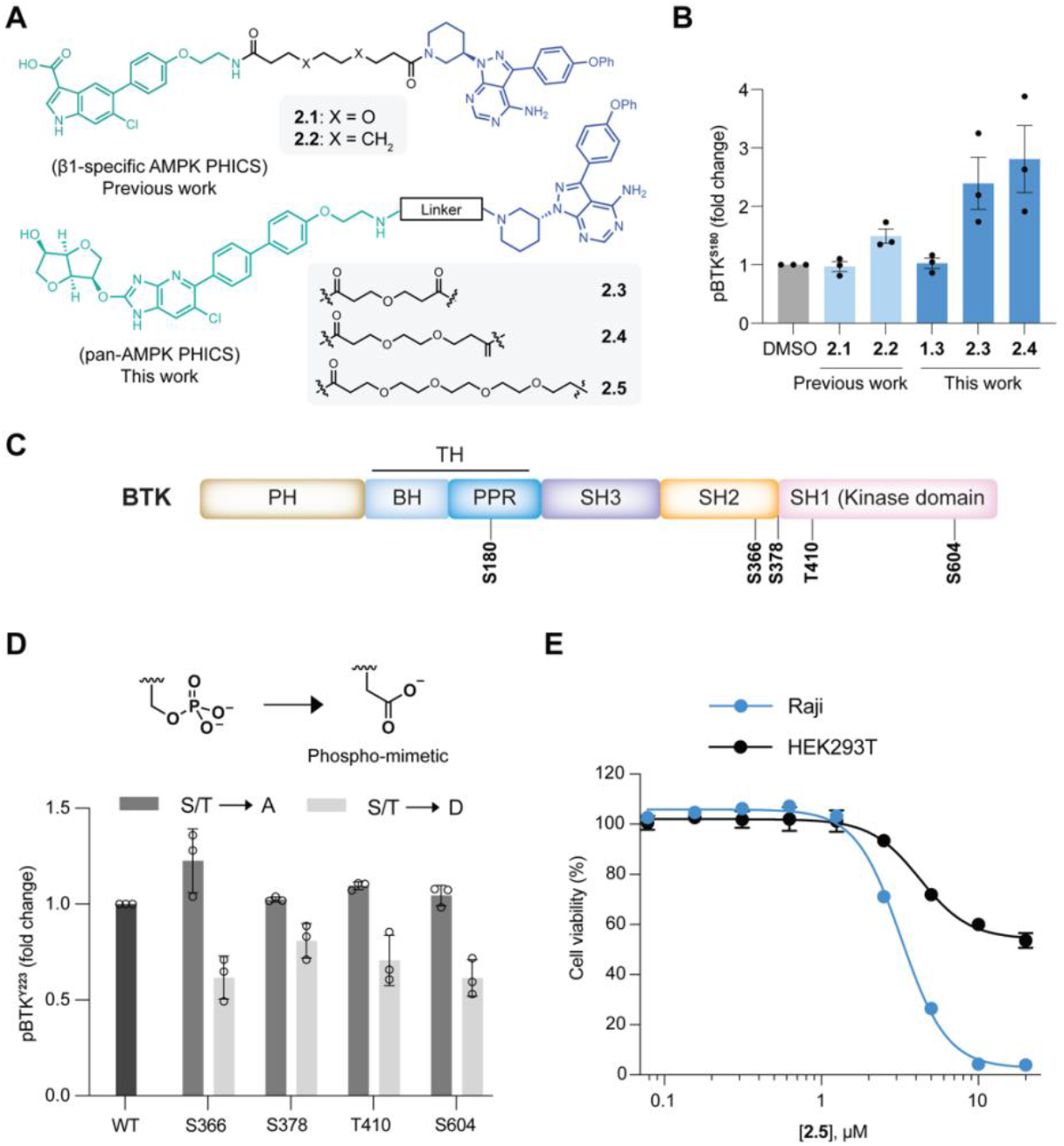
Development and functional characterization of BTK PHICS. (**A**) Chemical structures of BTK PHICS with varying linkers and chemotypes. (**B**) Induction of BTK^**S180**^ phosphorylation expressed in HEK293T cells treated with 5 µM of PHICS in serum-containing media. Results represent mean ± sd, relative pBTK^**S180**^ levels (fold-change with DMSO control). (**C**) Schematic representation of BTK domains (PH: Pleckstrin Homology; TH: Tec Homology; BH: BTK Homology region; PPR: PolyProline Region; SH3: Src-Homology 3; SH2: Src-Homology 2; Sh1: Src-Homology 1(Kinase domain). (**D**) Quantification of the effect of phospho-resistant and phospho-mimetic variants of BTK on its kinase activity measured by the autophosphorylation (pY223) levels. (**E**) CellTiter-Glo viability assay demonstrating the dose-dependent effects of PHICS **2.5** on Raji (EC_*50*_ = 3.2µM), and HEK293T cells viability.

### BTK PHICS induces death of ibrutinib-resistant cells

S180 phosphorylation is known to inhibit BTK activity^26^ and we, indeed, observed a decrease in BTK autophosphorylation at pY223 (a metric for kinase activity) in PHICS-treated cells (**Figure S5D**). The aforementioned studies focused on S180, and we were interested in determining if other sites were being phosphorylated. We treated HEK293T cells transiently expressing BTK with PHICS and performed mass spectrometry on immunoprecipitated BTK using DMSO-treated cells as a control. We used a previously reported method^27^ to determine the relative phosphorylation levels in PHICS and DMSO-treated cells and observed increased phosphorylation at S366, S378, T410, and S604 but not at T191 or T316 (**Figure S5E-J, S6**). T410 is located near the ATP binding pocket, while S366 and S378 are on the loops of the SH2 domain, which is critical for BTK activity. To determine the impact of these phosphorylations on BTK activity, we generated phospho-mimetic (Ser/Thr→Asp) and phospho-resistant (Ser/Thr→Ala) BTK variants at these sites. We assessed the activity of these variants by measuring the Y223 autophosphorylation and observed that the phospho-mimetic variants at S366, S378, T410, or S604 had reduced Y223 autophosphorylation (**Figure 2D, S5L**), but the control mutations (Ser→Ala) at these sites did not reduce Y223 phosphorylation. These studies suggest that inhibition of BTK activity by PHICS could arise via multiple mechanisms in addition to induction of S180 phosphorylation.

Motivated by the new inhibitory mechanisms of PHICS, we explored PHICS activity in the context of drug-resistant cancer cells. The FDA-approved BTK inhibitor, ibrutinib, prevents B-cell proliferation by blocking the aberrant BTK activity in B-cell cancers, such as mantle cell lymphoma (MCL) and chronic lymphocytic leukemia (CLL)^28^ but the emergence of ibrutinib resistance is a challenge.^29^ We hypothesize that despite utilizing the same BTK binder (ibrutinib), chimeric molecules can circumvent resistance arising from binding pocket mutations, owing to their event-driven mechanism of action rather than the conventional occupancy-driven approach.^30^ To test this hypothesis, we tested PHICS in reported^31^ BTK cell lines (Raji and Mino) against which Ibrutinib exhibits weak antiproliferative activity (EC_50_ = 14.5 µM for Raji and 8.4 μM for Mino cells). Strikingly, **2.5** inhibited the proliferation of Raji and Mino cells but not HEK293T cells (BTK negative cells), with an EC_50_ of 3.2 µM and 5.0 µM, respectively (**Figure 2E, S5M**). Since the dose-response curve’s baselines differ between cell lines, we computed area under the curve (AUC) and observed similar selectivity of **2.5** for BTK cell lines over HEK293T cells (**Figure S5N**).^32^ We note that the tested PHICS still exhibit moderate potency (low micromolar), and further structure-activity optimization will be needed for deep mechanistic investigations, an avenue we will pursue in the future.

### PHICS induces Liprin-α3 phase separation

Charged PTMs, like phosphorylation, play an essential role in regulating the formation and function of protein condensates that have a crucial role in processes such as metabolism,^33^ signaling complexes,^34^ gene transcription and translation,^35^ and synaptic transmission.^36^ Phosphorylation-induced phase separation of Liprin-α3, a component of specialized presynaptic structures, is responsible for regulating synaptic vesicle exocytosis (**Figure 3A)**.^20^ Since small-molecule binders of Liprin-α3 are unavailable, we used a chemogenetic tag (FKBP12^F36V^)^37^ fused to Liprin-α3 for recruiting AMPK to Liprin-α3 using PHICS. We synthesized several bifunctional molecules using pan-AMPK binder and FKBP12^F36V^ binder (AP1867) connected by linkers of various length and chemotype **(Figure 3B)**. In parallel, we generated HEK293T cell lines stably expressing mVenus–Liprin-α3 fused to FKBP12^F36V^ at either N- or C-termini **(Figure 3C)**. A two-dimensional screening of Liprin fusions and bifunctional molecules using automated imaging showed that only the C-terminal FKBP fusion (hereafter referred to as mVenus–Liprin-α3–FKBP12^F36V^) exhibited phase separation **(Figure 3D)**. Notably, neither the individual binders (MK8722 or AP1867) nor their combination exhibited phase separation. PHICS exhibited linker length dependence and PHICS treatment did not induce phase separation in cells expressing mVenus–Liprin-α3 that lacks FKBP12^F36V^ chemogenetic tag **(Figure 3D)**.

**Figure 3.**
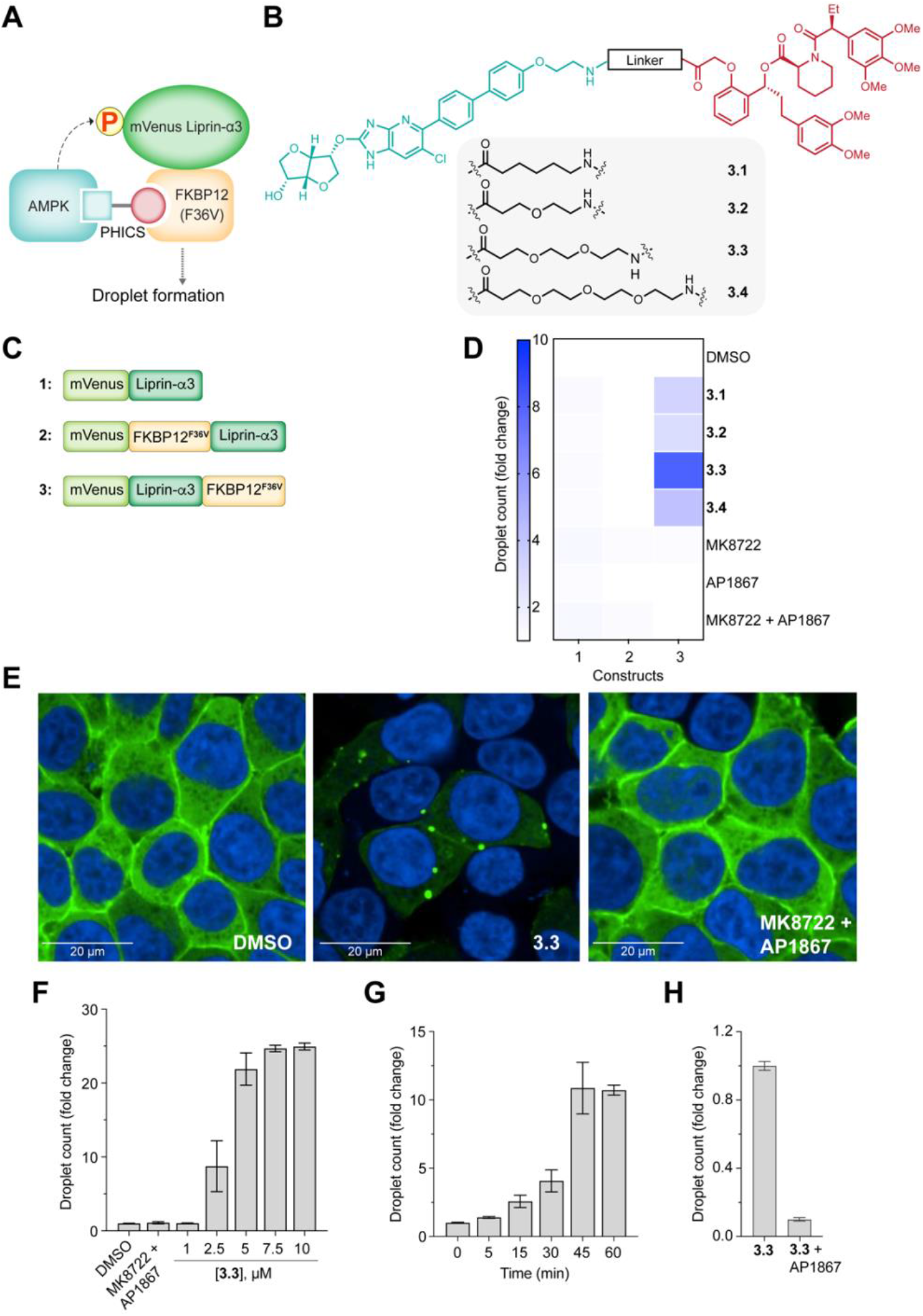
Phase separation of Liprin-α3. (**A**) Schematic representation of Liprin-α3–PHICS–AMPK ternary complex. (**B**) Chemical structures of Liprin-α3 PHICS with varying linker lengths and chemotypes. (**C**) Schematic representation of Liprin-α3 constructs showing N- or C-terminal fusion of FKBP12^F36V^. (**D**) Screening of Liprin-α3 constructs for the formation of Liprin-α3 droplets in the presence of PHICS (5 µM) or controls. (**E**) Confocal microscopic images (63x) of Liprin-α3 droplets in HEK293T cells stably expressing mVenus–Liprin-α3–FKBP12^F36V^. (**F, G**) Quantification of dose and time-dependent formation of Liprin-α3 droplets induced by **3.3**. (**H**) Competition experiment with FKBP12^F36V^ binder AP1867 (5 µM) with **3.3** (5 µM).

Among the PHICS tested compound **3.3** exhibited the most pronounced induction of Liprin-α3 phase separation, with robust droplet formation observed across a 1–10 µM concentration range **(Figures 3E, F, S7A)**. Importantly, neither individual binders (MK8722 or AP1867) alone nor their equimolar combination **(Figure S7A)** induced phase separation, reinforcing the necessity of optimizing linker length and chemotype to achieve the appropriate kinase–substrate proximity and orientation for phase separation. Additionally, the phase separation process induced by **3.3** was rapid, with detectable droplet formation within 15 minutes of treatment and saturation observed at approximately 45 minutes **(Figure 3G)**. Competition experiment with co-treatment with AP1867 abrogated PHICS activity, confirming that **3.3** triggered droplet formation depends on its bifunctional nature **(Figure 3H)**.

### Identification of phosphorylation sites responsible for Liprin-α3 droplet formation

Following confirmation of the bifunctional mechanism, we performed global phosphoproteomics to evaluate the selectivity of **3.3** and to identify the phosphosite responsible for PHICS-induced phase separation. Cells treated with **3.3** had similar global protein expression profile as cells treated with either DMSO (**Figure S7B**) or equimolar combination of MK87222 and AP1867 (**Figure S7C**). Furthermore, out of ≈42,000 identified phosphopeptides, only 17 phosphopeptides showed significant increase (fold-change >2.0, *p* < 0.05) in **3.3**-treated samples (**Figure 4A**). Of these 17 phosphopeptides, 14 are on mVenus-Liprin-α3 fusion, 2 on CASK (a Liprin-α3 interactor), and 1 on ELF4G, pointing to the exquisite specificity of PHICS. Finally, MK8722 treated cells showed an increase in phosphopeptides from known AMPK substrates^38^ (fold-change >2.0, *p* <0.05), but not on Liprin-α3 (**Figure S7D**), further suggesting that **3.3** is operating via bifunctional mechanism.

**Figure 4.**
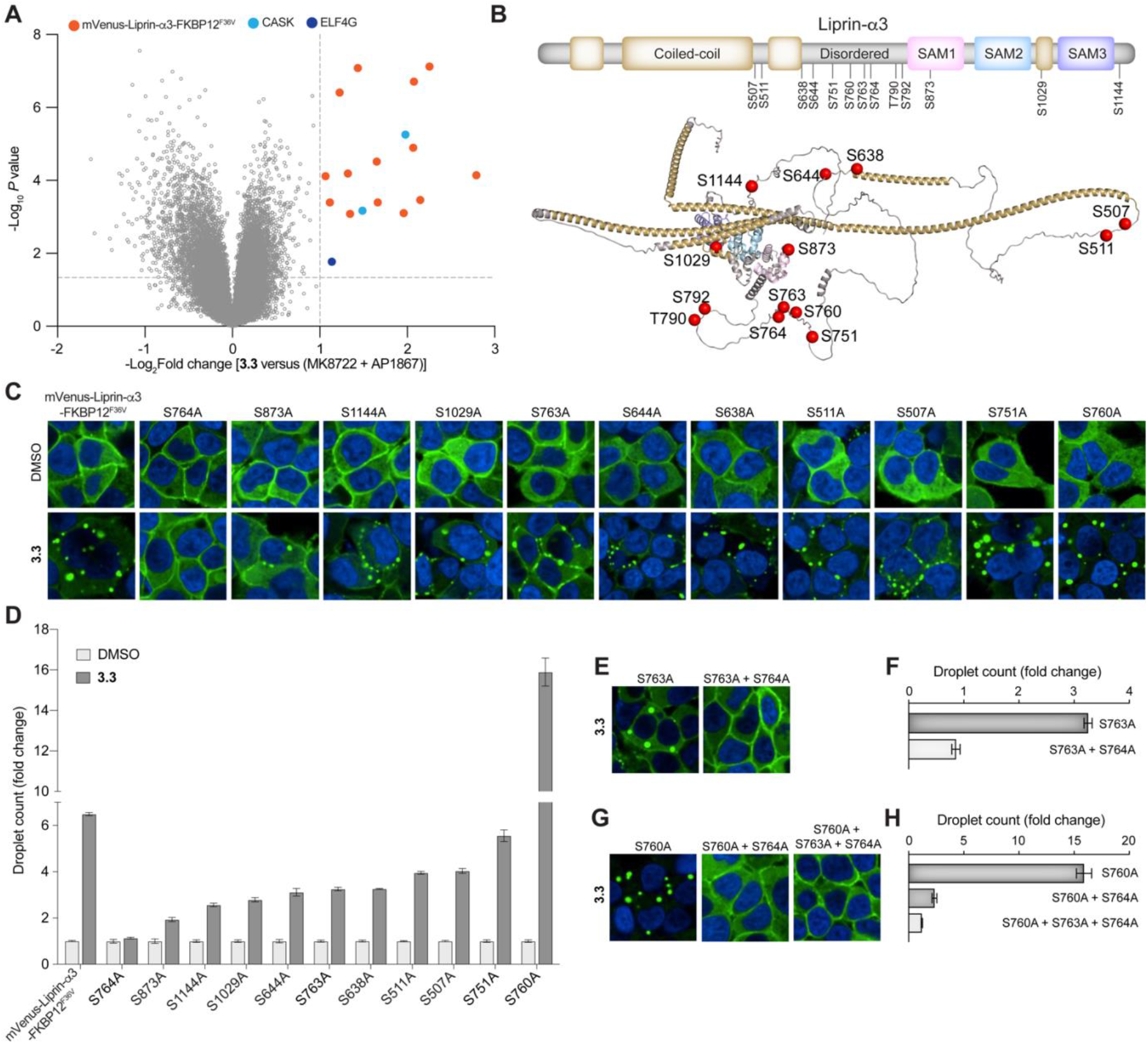
Global phosphoproteomics and validation of phosphosites responsible for Liprin-α3 droplet formation. (**A**) Volcano plots displaying the −log_10_ (*P*-value) versus log_2_ fold-change of phosphopeptides for **3.3** *vs*. combination of MK8722 and AP1867. (**B**) Schematic representation and alphafold2 structure of Liprin-α3 marked with potential phosphosites (red). Coiled-coil regions and sterile alpha motifs (SAM) are also highlighted in the schematic. (**C**) Confocal microscopic images (40x) of HEK293T stable cells expressing single alanine variant (i.e., S/T→A) of mVenus–Liprin-α3– FKBP12^F36V^. (**D**) Screening and validation of **3.3**-induced Liprin-α3 droplets in HEK293T stable cell lines expressing single alanine variants of mVenus–Liprin-α3–FKBP12^F36V^. (**E-H**) Confocal microscopic images (E, G) and quantification (F, H) of **3.3**-induced Liprin-α3 droplets in HEK293T stable cell lines expressing double and multiple alanine variants of mVenus–Liprin-α3–FKBP12^F36V^.

While global phosphoproteomics identified several **3.3** induced phosphosites that mostly reside in the disordered regions of the Liprin-α3, we did not observe phosphorylation at S760 that has been shown to trigger phase separation.^20^ The S760 is a basophilic site (Ser/Thr are surrounded by Arg/Lys), which complicates global phosphoproteomics studies as the digestion enzymes used (Trypsin/Lys-C) also cleave at Arg/Lys. Furthermore, AMPK has a higher propensity to phosphorylate basophilic site. Hence, we created an exhaustive list of potential phosphorylation sites considering not just the aforementioned phosphoproteomics data but also potential AMPK phosphosites^38, 40, 41^ and literature reports (**Figure 4B**).^39, 42, 43^ To validate these sites, we created 14 cell lines stably expressing the alanine variant of the phosphosite (i.e., S/T→A) (**Figure 4B, S7D)**. Of all the variants tested, S764A variant showed the most resistance to **3.3**-mediated droplet formation (**Figure 4C, D**). Interestingly, S760A or S763A variants alone were not resistant (**Figure 4C, D**), but composite mutations with S764A (i.e., double mutants S760A/S764A or S763A/S764A, or triple mutant S764A/S763A/S760A) were resistant (**Figure 4E-H**), further validating that S764 is the target phosphosite. Finally, the S760A variant showed two-fold higher propensity to form droplets than wild-type in presence of PHICS, opening avenues to examine the role of enhanced droplet formation on neuronal function. Overall, these results suggest that PHICS can redirect AMPK to phosphorylate S764 and induce phase-separation of Liprin-α3.

## Conclusions

We addressed issues with the previous PHICS platform that limited its applicability by designing, synthesizing, and validating the PHICS platform that recruits AMPK to target proteins. To do this, we used a pan-AMPK binder to generate AMPK PHICS for different target proteins and demonstrated their recruitment of multiple types of AMPK complexes. The AMPK PHICS can induce phosphorylation without serum starvation of cells and has demonstrated phosphorylation of an endogenous protein without the need for target overexpression. Application of AMPK PHICS to biologically relevant models revealed its potent inhibitory effects on cell proliferation in ibrutinib-resistant cells. Finally, further investigations using AMPK PHICS demonstrated its ability to initiate Liprin-α3 phosphorylation-mediated phase separation. Overall, these findings suggest that PHICS can be designed to induce proximity and the correct orientation of the kinase and the target protein for effective phosphorylation, emphasizing the potential utility of this bifunctional system in inducing biologically relevant phosphorylations on target proteins. The ability of these bifunctional compounds to modify the phosphorylation of native and disease-relevant proteins has therapeutic implications in health and disease.

## Acknowledgments

This work was supported by the Merkin Institute of Transformative Technologies in Healthcare, The Mark Foundation for Cancer Research, DARPA (HR00112120010), and NIH (1U01DK137242, R01GM137606, R01DK129464, R01GM132825). S.L. was supported by the NSF Graduate Research Fellowship (DGE-1745303). We thank Prof. Pascal Kaeser for providing Liprin plasmids.

## Author Contributions

R.P., S.V., P.K., S.L., P.K.T., and S.S. contributed equally to this paper.

