## Supporting Information for "Phosphorylation-inducing molecules for regulating dynamic cellular processes"

**Table of Contents**

1. Materials and methods. .......... S1

1. 1. *In vitro* kinase assay and western blotting to confirm the AMPK-PHICS-induced BRD4 phosphorylation .. S**Error! Bookmark not defined.**

1. 2. Cell culture……….......S2

1. 3. Western blotting to confirm the AMPK-PHICS-induced endogenous BCR-ABL phosphorylation. .......... S2

1. 4. Western blotting to confirm the AMPK-PHICS-induced BTK phosphorylation .......... S3

1. 5. Mass spectrometry analysis of the AMPK-PHICS-induced BTK phosphorylation. .......... S3

1. 6. Generation of phosphomimetic BTK variants and analysis of their effect on kinase activity by western blotting.S4

1. 7. Cell viability assay. .......... S4

1. 8. Generation of Liprin-α3 lentiviral plasmid constructs, stably expressing HEK293T cells and imaging, screening, and analysis. .......... S4

1.9. Global proteomics and phosphoproteomics……….... S5

*2.* Supplemental figures. .......... S6

### 1. Materials and methods

#### 1.1. *In vitro* kinase assay and western blotting to confirm the AMPK-PHICS-induced BRD4 phosphorylation.

The *in vitro* kinase reaction was performed in the presence of 20 nM AMPK (Complexes 1, 3, 4, and 5 from Invitrogen/Life Technologies, #PV4673, #PV6241, #A30485, #A30486; and Complex 2 from BPS Biosciences, #40703), 700 nM of truncated BRD4-GST (BPS Biosciences, #31044), 150 µM ATP, and 2.5 µM PHICS in 1X Kinase Buffer (Cell Signaling Technology, #9802) for 2 hours at room temperature. After treatment, the kinase reaction was quenched with SDS sample buffer by heating at 95 ˚C for 5 min. The samples were then resolved by 4-12% NuPAGE™ gel (#NP0336BOX) and transferred to the nitrocellulose (NC) membrane (#IB23002) using an iBlot™ 2 gel transfer device. After transfer, the NC membrane was incubated with a blocking buffer (5% BSA in 1xTBS and 0.1% Tween 20) for 1 hour at room temperature. The membrane was then incubated overnight on a shaker at 4 ˚C with primary antibodies: anti-phospho-AMPK-motif antibody (Cell Signaling Technology, #5759) and anti-GST antibody (Cell Signaling Technology, #2624) at 1:1000 dilutions were prepared in the blocking buffer. Following overnight incubation, the unbound primary antibodies were washed out three times with the blocking buffer at 10-15 min of each wash. The membrane was incubated further with secondary antibodies: Donkey anti-rabbit antibody (LI-COR Biosciences, #926-68073) and anti-mouse antibody (LI-COR Biosciences, #926-32212) prepared in the blocking buffer at 1:10,000 dilutions. The unbound secondary antibodies were washed out and finally rinsed with deionized water. The blots were developed using the LI-COR Odyssey CLx Imaging System.

#### 1.2. Cell culture

All cell lines used in this study were purchased from the American Type Culture Collection (ATCC) and maintained in 37 °C, 5% $CO_2$ humidified incubators. All cell culture media and reagents were purchased from Gibco. HEK293T (CRL-3216) cells were cultured in Dulbecco's Modified Eagle's Medium (DMEM) with 1% sodium pyruvate, 10% fetal bovine serum, and 1% penicillin-streptomycin. Raji (CCL-86), Mino (CRL-3000), and K-562 (CCL-243) cells were cultured in RPMI-1640 and 10% fetal bovine serum and 1% penicillin-streptomycin.

#### 1.3. Western blotting to confirm the Gen 2 AMPK-PHICS-induced endogenous BCR-ABL phosphorylation.

The PHICS compound solutions (for dose and competition with MK8722) were prepared at the final concentration of 0.1% DMSO in RPMI complete media. K562 cells cultured in RPMI media were counted, and $10x10^6$ cells per condition were taken and incubated with PHICS solutions at the density of $1x10^6$ cells per mL for 4 hours. After compound treatment, cells were washed with ice-cold PBS, collected as a pellet, and stored at -80 ˚C until further use. The cell pellet was then treated (with gentle agitation/rotation) for one 1 hour with M-PER (Mammalian Protein Extraction Reagent, ThermoFisher, #78501) supplemented with 1X cOmplete EDTA-free Protease Inhibitor (Roche, #4693159001) and 1X PhosSTOP Phosphatase Inhibitor (Roche, #4906837001). The cell lysates were centrifuged at 16,000 x g for 10 minutes at 4 ˚C, and the resulting supernatant was collected in clean tubes to estimate the total protein concentration with Pierce BCA Protein Assay Kit (ThermoFisher, #23225). Immunoprecipitation (IP) of the BCR-ABL protein was performed using the Dynabeads Protein G (ThermoFisher, #10004D) according to manufacturer instructions, using the ABL primary antibody (Cell Signaling Technology, #2862) at 1:1000 (1µg AbL Ab / 1000µg total lysate) concentration and overnight protein binding at 4 °C on a rotator. All protein-bead samples were eluted with 50 µL 4X Sample buffer (BioRad, #1610747) with ß-ME added freshly, and heated at $95^0$C for 5 minutes to obtain IP eluates. Equal volumes of IP eluate were resolved by SDS-PAGE, transferred to nitrocellulose (NC) membranes using iBlot™ 2 with transfer stacks (ThermoFisher, #IB401002), followed by blocking with blocking buffer (5% BSA in 1xTBST) at room temperature for 1 hour and rinsed with 1x TBST. Membranes were incubated overnight at 4 °C with primary antibodies, anti-phospho-AMPK-motif antibody (Cell Signaling Technology, #5759), and mouse anti-ABL (24-11) mAb #SC-23 (Santa Cruz Biotechnology) at 1:1000 and 1:200 dilutions respectively. Following overnight incubation, the membranes were then washed with 1x TBST for 3 x 5 minutes and incubated with 1:10,000 LI-COR IRDye® secondary antibodies (680RD donkey anti-rabbit #926-68073, and 800CW donkey anti-mouse #926-32212) for 1 hour at room temperature. After washing again and rinsing with deionized water, the membranes were imaged on an Odyssey imager (LI-COR).

#### 1.4. Western blotting to confirm the AMPK-PHICS-induced BTK phosphorylation.

BTK-FLAG plasmid (clone ID: OHu20257) was purchased from GenScript. HEK293T cells were seeded in twelve-well cell culture plates (Corning) and grown for 24 hours to reach 70-80% confluency. The next day, BTK transfection was performed using TransIT-LT1 transfection reagent (Mirus Bio, #MIR2306) according to manufacturer instructions and incubated for 24 hours. Following incubation, PHICS compound solutions were prepared at the final concentration of 0.1% DMSO in DMEM media (with and without serum) and added to the cells. The cells were then incubated for 4 hours, washed out compounds with ice cold PBS, and cells with the entire plate were stored at -80 ˚C until further use. The preparation of cell lysates and subsequent western blotting procedure was the same as detailed in section 1.3. (see BCR-ABL sample preparation and western blotting). An equal amount of total lysate (20µg) was used for western blotting. The primary antibodies used were mouse anti-phospho-BTK$^{S180}$ mAb (Cell Signaling Technology, #3537) and rabbit anti-DYKDDDDK-Tag mAb (Cell Signaling Technology, #14793) at 1:1000 dilutions.

#### 1.5. Mass spectrometry analysis AMPK-PHICS induced BTK phosphorylation.

The preparation of cell lysates after BTK-FLAG transfection into HEK293T cells and subsequent compound treatment was the same as detailed in section 1.4. The BTK was then immunoprecipitated using anti-FLAG M2 magnetic beads (ThermoFisher, #A36798) according to manufacturer instructions. Briefly, an equal amount of total lysate was incubated with anti-FLAG beads for protein binding at 4 ˚C on a rotator. The samples were eluted with 150 µL 1X sample buffer (BioRad, #1610747; 4X was diluted to 1X) with ß-ME added freshly and heated at 95$^0$C for 5 minutes to obtain IP eluates. Equal volumes of IP eluate were resolved by SDS-NuPAGE gel. The resolved gels were Coomassie-stained to visualize and then extract the appropriate protein bands by cutting. The samples were submitted to the Taplin Mass Spectrometry Facility at Harvard Medical School to identify the induced phospho-sites after in-gel trypsinization. **LC-MS/MS data process and analysis:** The raw files were analyzed using Proteome Discoverer (PD) 2.5 software package (Thermo Fisher). Sequest HT was used to search against human UniprotKB/Swiss-Prot Homo sapiens proteome database (*www.uniprot.org*). Minora Feature Detector was also included in the processing step. Maximum missed cleavage was set to 2 and minimum peptide length was set to 6. Precursor mass tolerance was set to 10 ppm and fragment mass tolerance was set to 0.6 Da, respectively. Methionine oxidation (15.995 Da) and serine/threonine phosphorylation (79.966Da) were set as dynamic modifications. Both the maximum dynamic and equal modification per peptide were set to 4. N-terminal glutamate cyclization were set as peptide dynamic modification. Acetylation was set as protein terminus dynamic modification. Cysteine alkylation was set as static modification. Target Decoy PSM Validator was used for PSM validation with FDR set to be 0.01. IMP-ptmRS was added after Target Decoy PSM to calculate the probability of phosphorylation on detected amino acid. Tyrosine-protein kinase BTK (Q06187) was detected as the most abundant protein. The peptide groups identified from BTK were filtered with the following parameters in PSM: ptmRS phospho site probabilities = 100, XCorr > 1.5 to locate Ser/Thr that were modified by AMPK. To compare the effect of PHICS treatment against DMSO, the phosphorylated and unphosphorylated peptide abundance were used to calculate the approximate phosphorylation percentage using equation % Phosphorylation = (Abundance of phosphorylated peptide/ Abundance of phosphorylated peptide + unphosphorylated peptide). The result was exported and replotted in GraphPad Prism 9.0.

#### 1.6. Generation of phosphomimic BTK variants and analysis of their effect on kinase activity by western blotting.

After identifying the sites of PHICS-induced phosphorylation on BTK, site-directed mutagenesis (SMD) was used to generate phosphomimic BTK variants. The variants were then transfected into HEK293T cells, incubated for 48 hours, and checked for their effect on BTK kinase activity by measuring pY223 levels on western blots. The preparation of cell lysates and subsequent western blotting procedure was the same as detailed in section 1.4. The primary antibodies used were rabbit anti-phospho-BTK$^{Y223}$ mAb (Cell Signaling Technology, #87141) and mouse anti-DYKDDDDK-Tag mAb (Cell Signaling Technology, #8146) at 1:1000 dilutions.

#### 1.7. Cell viability assay

Compounds were dispensed at varying programmed concentrations to white opaque 384-well plates (Corning, #8867BC) using a D300e digital dispenser (Tecan) and T8+ dispensehead cassettes (#30097370). All cell lines (2000 cells per well) were seeded at 40µL onto the compound-containing plates and gently pipetted up and down to mix. The plates were then incubated at 37°C, 5% $CO_2$, for 72 hours. After incubation, the plates were taken out and equilibrated to room temperature. CellTiter-Glo reagent (Promega, #G7572) was thawed and diluted at a 1:1 ratio with PBS buffer, used according to manufacturer instructions, and the luminescence signal was read using EnVision plate reader (PerkinElmer). Data were analyzed using Prism 9 (GraphPad) software. **Calculation of area under the curve (AUC):** The dose-response curves for the cell line were fitted using a nonlinear regression method ("Inhibitor vs. Response – Variable Slope (four parameters)") in GraphPad Prism 10.4. The fitted parameters (Bottom, Top, EC50, Hill slope) were subsequently imported into MATLAB to generate the fitted curves. The minimum and maximum concentrations were used as the lower and upper bounds for the x-values, respectively. The area under the curve (AUC) was then calculated in MATLAB using Simpson's integration method, as described by the following equation:

$$\mathrm{AUC} = \frac{h}{3}\left(Y(1) + 4\sum Y(2:2:end-1) + 2\sum Y(3:2:end-2) + Y(end)\right)$$

where,

*h* is the step size

*Y(1)* is the first value in Y array, corresponding to the function value at $X_{min}$.

Y(end) is the last value in Y array, corresponding to the function value at $X_{max}$.

Y(2:2:end-1) : define odd indexed function values.

Y(3:2:end-2) : define even indexed function values.

#### 1.8. Generation of Liprin-α3 lentiviral plasmid constructs, stably expressing HEK293T cells, and Imaging, screening, and Analysis.

Lentiviral vector generation: gBlocks for Liprin-α3, its variants, $FKBP12^{F36V}$, and mVenus were designed and cloned into a lentivector using Gibson ligation. Stable line generation: Viral packaging plasmids psPAX2 (Addgene, #12260) and pMD2.G (Addgene, #12259) together with lentiviral Liprin-α3 plasmids were transfected to 293T cells in a 2:1:3 ratio using TransIT®-LT1 transfection reagent (MirusBio, #MIR2300) following the manufacturer's guidelines. Lentiviruses were collected 24 and 48 h after transfection and filtered with 0.45-µm Millex®-HP PES filters (Millipore, #SLHPM33RS). For transduction in HEK293T cells, the viral supernatant was mixed with culture media in a 1:1 ratio with 10 µg/ml Polybrene transfection reagent (Millipore Sigma, #TR-1003-G), then selected with 2 µg/mL puromycin after at least 24 h of transduction. Finally, antibiotic-selected cells were maintained in 1 µg/mL puromycin for routine culture. Imaging Screening and Analysis: HEK293T cells stably expressing Liprin-α3 constructs were seeded in 100 µl of 40,000 cells per well into PhenoPlate™ 96-well microplates (PerkinElmer, # 6055302). 24 hours after attachment, compound dilutions were directly printed into cell-seeded plates using Tecan D300e Digital Dispenser (Tecan) and HP T8+ dispense head cassettes (#F0L59A). The plates were incubated for 0-1 hour (for time-dependent experiments, single and multiple analine variants screening) and 4 hours (for single- and multiple-dose experiments). After 0-4 hours of incubation, the 96-well plates were washed once with PBS, then fixed in 4% paraformaldehyde (15 minutes) and stained by Hoechst 33342 (ThermoFisher Scientific, #H3570). High-content confocal imaging of cells was performed for each plate using an Opera Phenix imaging system, followed by automated analysis using Harmony Software v4.9 or PhenoLogic 5.0 (PerkinElmer/Revvity) analysis software. The number of mVenus droplets were identified using the software's automated spot-based analysis pipeline, which normalized the spot number per cell in DMSO-treated wells to determine the droplet fold change of each compound across doses. For each condition at least 30,000 cells were analyzed and these normalized values were used to generate a heatmap or bar graphs using GraphPad Prism 9.

#### 1.9. Global proteomics and phosphoproteomics.

##### 1.9.1. AMPK-Liprin-α3 PHICS:

HEK293T cells stably expressing mVenus-Liprin-α3-$FKBP12^{F36V}$ were cultured in DMEM supplemented with 10% FBS, 1% Penstrep, and 2µg/ml puromycin. They were cultured in a T-75 flask to reach around 90% confluence (around $15x10^6$ cells) and were treated with the following compounds (in four replicates) for 1 hour: 1. DMSO (Vehicle control), 2. PHICS 13 (bifunctional) at 5 µM, 3. AMPK binder (A) at 5 µM, and 4. The combo of AMPK binder (A) and FKBP-binder) at 5 µM each at the final concentration of 0.15% DMSO. After treatment, the cells were washed with ice cold PBS, trypsinized, collected in 15ml tubes, and washed with ice cold PBS. The cell pellets were transferred to 1.5 epi tubes (~$15x10^6$ cells per tube) and stored at -80 ˚C until further use.

##### 1.9.2. Sample preparation for mass spectrometry to analyze global phopsho-proteomics:

Cell pellets were lysed by EasyPep Lysis Buffer (Thermo #A45735) with protease and phosphatase inhibitors (Halt Protease and PPase Inhibitor Mix, Thermo #78444). According to the manufacturer's instructions, protein concentration was determined with the BCA assay (Rapid Gold BCA Kit, Thermo #A53225). Next, samples were reduced with 5mM TCEP (Thermo #20490), alkylated with 10 mM iodoacetamide (Thermo # 122270250) that was quenched with 10 mM DTT

(Thermo #20290). A total of 250 µg of protein lysate was cleaned using SP3 resin beads (Hydrophilic and Hydrophobic, Thermo # 09-981-121 and 09-981-123) according to manufacturer's protocol. Next, Trypsin/Lys-C Mix (Promega # V5072) in 200 mM EPPS pH 8.5 (1:100 ratio) was added and samples digested for 16 h. To the liquid containing digested peptide, acetonitrile was added to have 30% acetonitrile in final solution and TMT reagents in (1:1 ratio of TMT reagent to peptide, TMTpro 18-plex Kit, Thermo #A52045) were added.

A small portion of TMT samples was pooled to check labeling efficiency >95%. TMT reaction was quenched with 50% hydroxylamine solution (Thermo #90115). For total proteomic analysis, ~100 µg of the pooled sample was cleaned using SP3 resin protocol, fractionated with Pierce High pH Reversed-phase Peptide Fractionation Kit (Thermo #84868) according to manufacturer's protocol and the resulting 10 fractions were subsequently analyzed by LC-MS/MS on an Orbitrap Eclipse Tribrid instrument.

For the phosphoproteomics, the rest of the samples were vacuum centrifuged to dryness and desalted using Peptide Desalting Spin Columns (Thermo #89851). Next, the High-Select $TiO_2$ Phosphopeptide Enrichment Kit (Thermo #32993) and High-Select Fe-NTA Phosphopeptide Enrichment Kit (Thermo #A32992) was used to enrich phosphopeptides according to SMOAC (Sequential enrichment from Metal Oxide Affinity Chromatography) protocol. Pooled samples were then combined and vacuum centrifuged to dryness. Pooled phosphopeptides were fractionated with basic pH reversed-phase (Thermo). The resulting 36 fractions were combined into 12 fractions and subsequently analyzed by LC-MS/MS on an Orbitrap Eclipse Tribrid instrument.

#### 1.9.3. LC-MS/MS analysis of digested proteins:

For LC-MS/MS analysis, samples were reconstituted in 2% ACN, 0.2% formic acid solution, sonicated, and centrifuged at 10,000g for 1 minute. The LCMS-vials (Thermo Fisher, cat. 6PK1655) containing supernatant were subsequently loaded onto a Vanquish Neo UHPLC system (Thermo Fisher) connected to an Orbitrap Eclipse mass spectrometer (Thermo Fisher) equipped with a NanoFlex ion source. For total proteomics samples, peptides were separated on a 60 cm column (IonOpticks, cat. AUR3-60075C18-TS) using a 130-minute gradient (0.0-1.0 min, 3%-5% ACN, 1.0-69.0 min, 5%-25% ACN, 69.0-89.0 min 25%-40% ACN, 89.0-104.0 min 40%-60% ACN, 104-120 min 95% ACN, wash) at a flow rate of 0.3 µL/min. MS1 data were acquired in Orbitrap mode with a resolution of 120,000, standard AGC target, and auto maximum injection time. Charge states from 2+ to 6+ were included, and a dynamic exclusion time of 30 seconds was used. MS2 scans were isolated with the quadrupole and fragmented using HCD with a fixed collision energy of 35% and a 0.7 m/z isolation window. The normalized AGC target for MS2 was set to 200%, and fragment ions were detected in the Orbitrap at a resolution of 50,000 with a defined first mass of m/z 120. For phosphoproteomic samples, peptides were separated on a 60 cm column (IonOpticks, cat. AUR3-60075C18-TS) using a 180-minute gradient (0.0-6.0 min, 2%-3% ACN, 6.0-121.0 min, 3%-25% ACN, 121.0-152.0 min 25%-40% ACN, 152.0-161.0 min 40%-60% ACN, 161-180 min 95% ACN, wash) at a flow rate of 0.3 µL/min. MS1 data were acquired in Orbitrap mode with a resolution of 120,000, standard AGC target, and auto maximum injection time. Charge states from 2+ to 6+ were included, and a dynamic exclusion time of 60 seconds was used. MS2 scans were isolated with the quadrupole and fragmented using HCD with a fixed collision energy of 35% and a 0.7 m/z isolation window. The normalized AGC target for MS2 was set to 200%, and fragment ions were detected in the Orbitrap at a resolution of 50,000 with a defined first mass of m/z 120.

##### 1.9.4. LC-MS/MS data processing and analysis

Proteome Discoverer 2.5 (Thermo Fisher) was used to process the raw data, utilizing Sequest HT for peptide identification against the human reference proteome (Uniprot proteome ID: UP000005640_9606). The search criteria allowed up to two missed cleavages and included peptides with at least 6 amino acids. Precursor mass tolerance was set at 10 ppm, and fragment mass tolerance was fixed at 0.02 Da. The dynamic modifications accounted for methionine oxidation (+15.995 Da), phosphorylation (+ 79.966 Da) on serines, threonines, and tyrosines. *N*-terminal dynamic protein modifications included – (Acetyl, +42.011 Da), (Met-loss, -131.040 Da), (Met-loss+Acetyl, -89.030 Da). Static modifications incorporated cysteine carbamidomethylation (+57.02146 Da), and TMT reagent addition (+304.207 Da) on peptide *N*-termini and lysines. Peptide-spectrum match (PSM) validation was performed using Percolator with a strict false discovery rate (FDR) of 0.01. IMP-ptmRS node was used for phosphosite localization. The resulting data was exported as CSV for importing into Perseus,[1] where reporter ion abundances were $log_2$-transformed and normalized by subtracting each column's median, replicates were grouped according to experimental conditions, and Welch's t-tests were performed to identify phosphopeptides with significantly altered abundances. Volcano plots were prepared in GraphPad Prism.

### 2. Supporting Figures

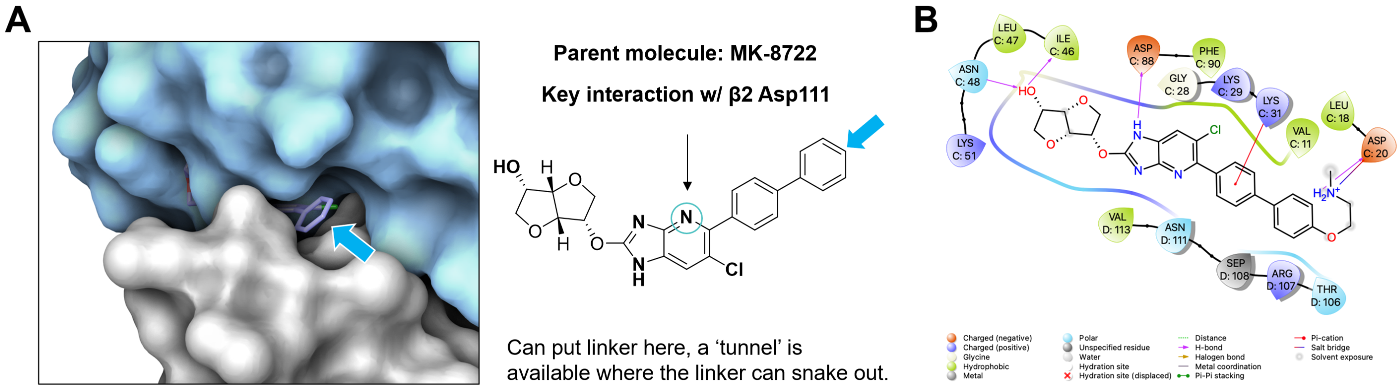


**Figure S1.** Docking of MK8722 and its analogues. Docking was performed with the standard precision protocol using Schrödinger Maestro v11.6. The ligands were prepared by generating possible states at pH 7.0 ± 2.0 using Epik, desalted, and subjected to OPLS3e force field. 2D ligand interaction maps were generated from the docking results to predict linker attachment sites. (A) Surface protein view showing buried molecule MK8722 between AMPK subunits. The Blue arrow represents the solvent-accessible location. (B) Ligand-residue interaction map between modified MK8722 analog and nearby amino acids.

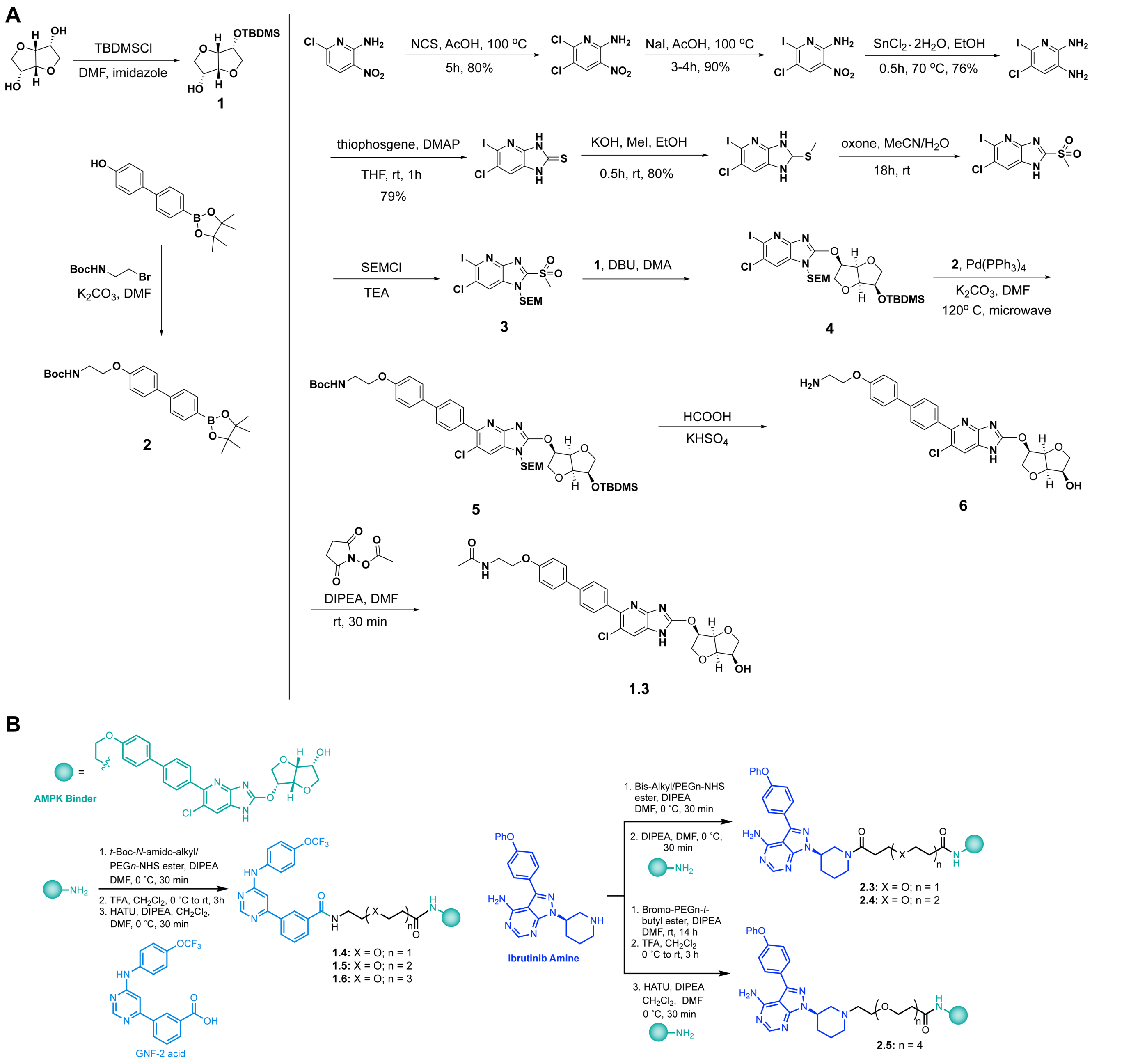


**Figure S2**. (**A**) Synthesis of the AMPK binder used for building various PHICS. (**B**) General synthetic scheme for generating AMPK-PHICS for multiple targets through amide coupling or *N*-alkylation varying linkers.

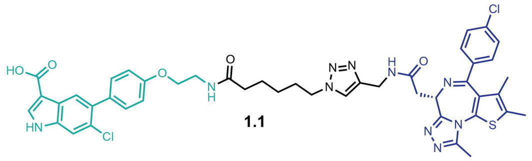

**1.1**

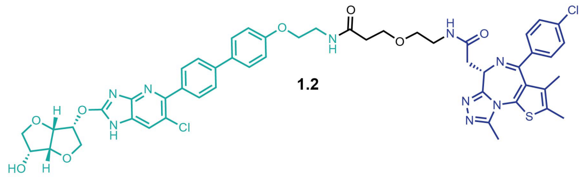

**1.2**

**Figure S3.** Structures of the AMPK-BRD4 PHICS used for the biochemical experiment to determine isoform specificity.

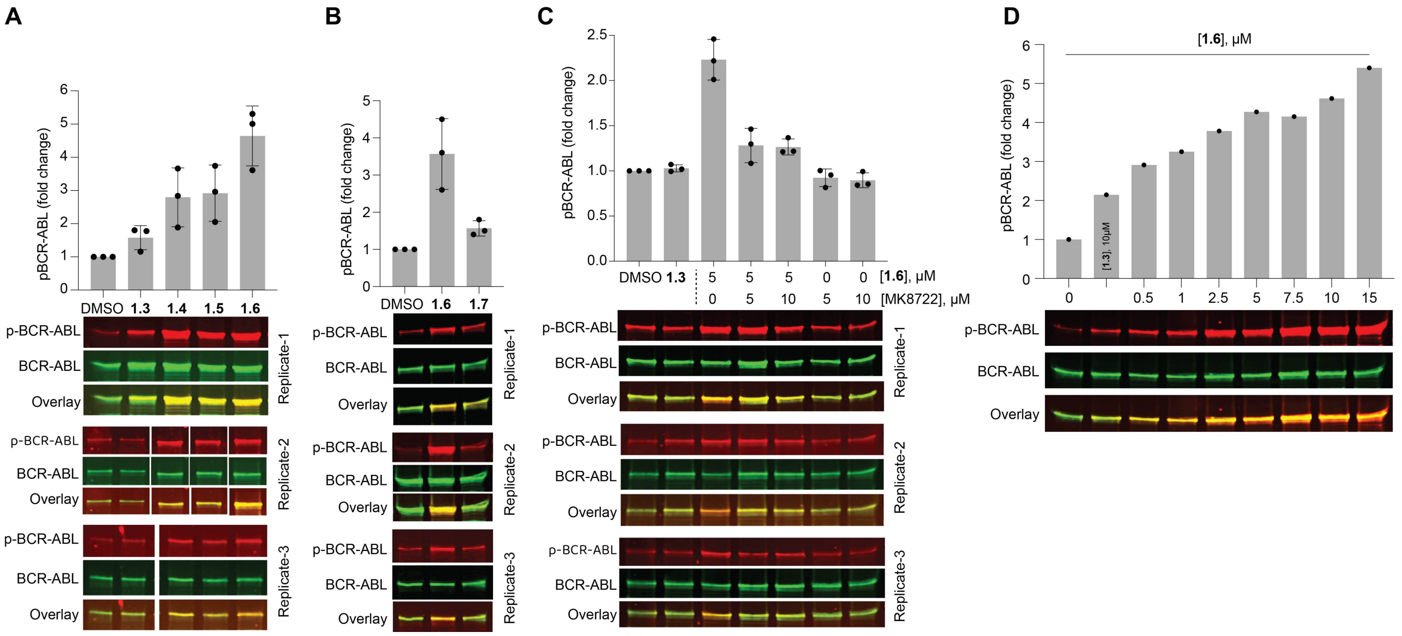


**Figure S4**. Functional assessment of AMPK-BCR-ABL PHICS. Panels (**A**, **B**, and **C**): Quantification of pBCR-ABL levels from three independent experiments represent panels (E, **F**, and G) of the figure 1. (**D**) Induction of dose-dependent phosphorylation of endogenous BCR-ABL by 1.**6** in K562 cells.

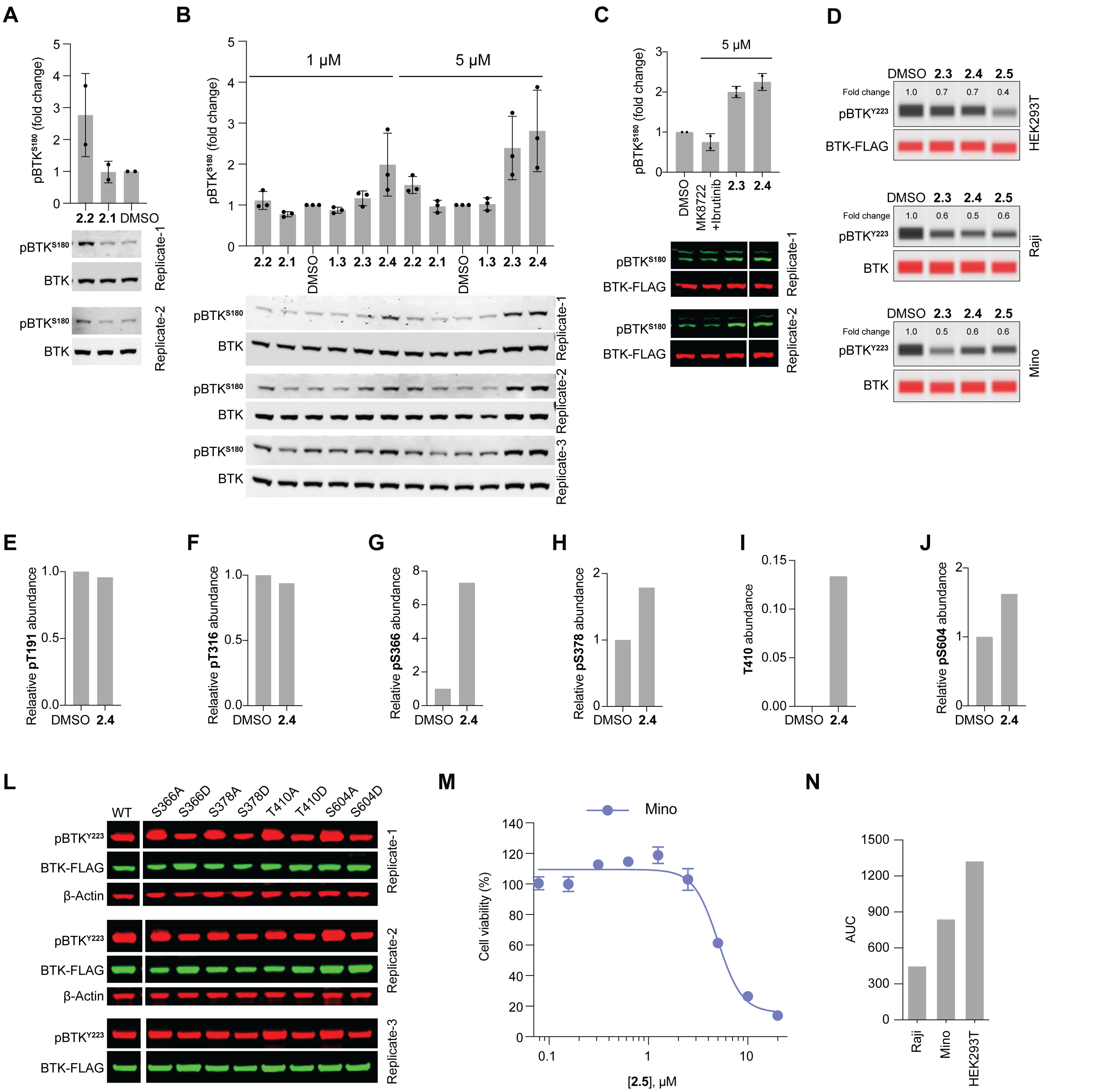


**Figure S5**. Functional assessment of AMPK-BTK PHICS. (**A**, **B**, and **C**) Induction of $BTK^{S180}$ phosphorylation by first-generation PHICS in serum free media (**A**) and second-generation PHICS in complete (serum containing) media (**B** and **C**). (**D**) Effect of PHICS on BTK kinase activity measured by BTK autophosphorylation (pY223) in HEK293T, Raji, and Mino cells. (**E-K**) Quantification of PHICS-induced phosphorylation of BTK sites T191 (**E**), T316 (**F**), S366 (**G**), S378 (**H**), T410 (**I**), and S604 (**J**) identified by mass spectrometry. (**L**) Western blots from three independent experiments for the effect of phosphomimic variants of BTK on its kinase activity measured by the autophosphorylation (pY223) levels supporting figure 2D. (**M**) CellTiter-Glo viability assay demonstrating the dose-dependent effects of PHICS **2.5** on Mino ($EC_{50}$ = 5.0 µM). (**N**) The area under the curve (AUC) of the dose-response curves for the indicated cell lines (Figure 2E) was calculated, to assess the relative potency and total level of inhibition observed for **2.5**, in MATLAB using Simpson's integration method, as described in the method section 1.7.

A

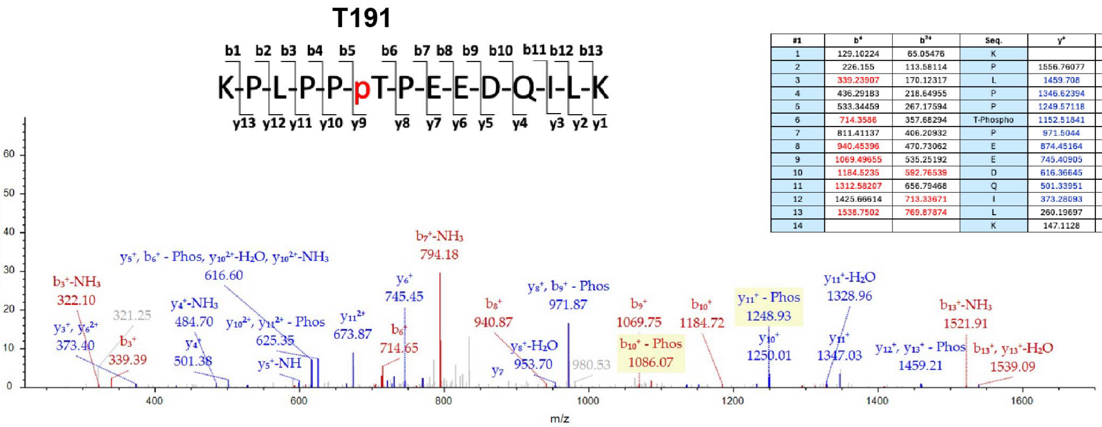


| #1 | b⁺ | b²⁺ | Seq. | y⁺ | y²⁺ | #2 |
|---|---|---|---|---|---|---|
| 1 | 129.10224 | 65.05476 | K | | | 14 |
| 2 | 226.155 | 113.58114 | P | 1556.76077 | 778.88402 | 13 |
| 3 | 339.23907 | 170.12317 | L | 1459.708 | 730.35764 | 12 |
| 4 | 436.29183 | 218.64955 | P | 1346.62394 | 673.81561 | 11 |
| 5 | 533.34459 | 267.17594 | P | 1249.57118 | 625.28923 | 10 |
| 6 | 714.3586 | 357.68294 | T-Phospho | 1152.51841 | 576.76284 | 9 |
| 7 | 811.41137 | 406.20932 | P | 971.5044 | 486.25584 | 8 |
| 8 | 940.45396 | 470.73062 | E | 874.45164 | 437.72946 | 7 |
| 9 | 1069.49655 | 535.25192 | E | 745.40905 | 373.20816 | 6 |
| 10 | 1184.5235 | 592.76539 | D | 616.36645 | 308.68686 | 5 |
| 11 | 1312.58207 | 656.79468 | Q | 501.33951 | 251.17339 | 4 |
| 12 | 1425.66614 | 713.33671 | I | 373.28093 | 187.1441 | 3 |
| 13 | 1538.7502 | 769.87874 | L | 260.19697 | 130.60207 | 2 |
| 14 | | | K | 147.1128 | 74.06004 | 1 |

B

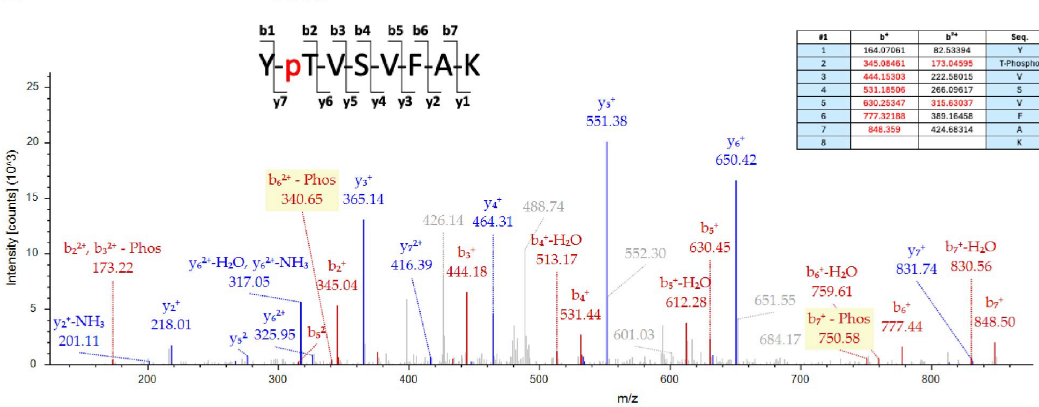


| #1 | b⁺ | b²⁺ | Seq. | y⁺ | y²⁺ | #2 |
|---|---|---|---|---|---|---|
| 1 | 164.07061 | 82.53394 | Y | | | 8 |
| 2 | 345.08461 | 173.04595 | T-Phospho | 831.4012 | 416.20424 | 7 |
| 3 | 444.15303 | 222.58015 | V | 650.38719 | 325.69723 | 6 |
| 4 | 531.18506 | 266.09617 | S | 551.31877 | 276.16033 | 5 |
| 5 | 630.25347 | 315.63037 | V | 464.28675 | 232.64701 | 4 |
| 6 | 777.32188 | 389.16458 | F | 365.21333 | 183.1128 | 3 |
| 7 | 848.359 | 424.68314 | A | 218.14992 | 109.5786 | 2 |
| 8 | | | K | 147.1128 | 74.06004 | 1 |

C

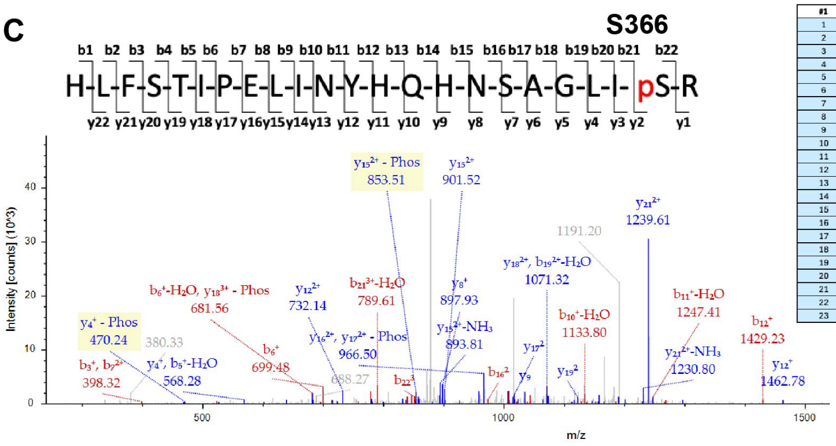


| #1 | b⁺ | b²⁺ | Seq. | y⁺ | y²⁺ | #2 |
|---|---|---|---|---|---|---|
| 1 | 138.06619 | 69.53673 | H | | | 23 |
| 2 | 251.15025 | 126.07876 | L | 2590.28136 | 1295.64432 | 22 |
| 3 | 398.21867 | 199.61279 | F | 2477.1973 | 1239.10229 | 21 |
| 4 | 485.25069 | 243.12899 | S | 2330.12888 | 1165.56808 | 20 |
| 5 | 586.29387 | 293.65282 | T | 2243.09687 | 1122.05227 | 19 |
| 6 | 699.38244 | 350.19486 | I | 2142.04919 | 1071.52623 | 18 |
| 7 | 796.4352 | 398.72124 | P | 2028.96511 | 1014.9862 | 17 |
| 8 | 925.47779 | 463.24254 | E | 1931.91235 | 966.45853 | 16 |
| 9 | 1038.56186 | 519.78457 | L | 1802.86976 | 901.93385 | 15 |
| 10 | 1151.64592 | 576.3266 | I | 1689.78569 | 845.39149 | 14 |
| 11 | 1265.68885 | 632.84443 | N | 1576.70163 | 788.84946 | 13 |
| 12 | 1428.75218 | 714.87936 | Y | 1462.6587 | 731.32935 | 12 |
| 13 | 1565.81079 | 783.40918 | H | 1299.59536 | 650.29443 | 11 |
| 14 | 1693.89667 | 847.43497 | Q | 1162.53675 | 581.76715 | 10 |
| 15 | 1830.95828 | 915.97939 | H | 1034.47789 | 517.23808 | 9 |
| 16 | 1944.97151 | 972.98939 | N | 897.41789 | 449.21372 | 8 |
| 17 | 2032.00353 | 1016.50541 | S | 783.37305 | 392.00735 | 7 |
| 18 | 2103.04065 | 1052.52052 | A | 703.34075 | 351.67302 | 6 |
| 19 | 2160.06211 | 1080.53426 | G | 632.3056 | 315.17032 | 5 |
| 20 | 2273.14618 | 1137.07673 | L | 568.28544 | 284.14405 | 4 |
| 21 | 2386.23024 | 1193.61876 | I | 452.21051 | 226.10525 | 3 |
| 22 | 2553.2286 | 1277.11794 | S-Phospho | 342.11731 | 171.56229 | 2 |
| 23 | | | R | 175.1119 | 88.06311 | 1 |

D

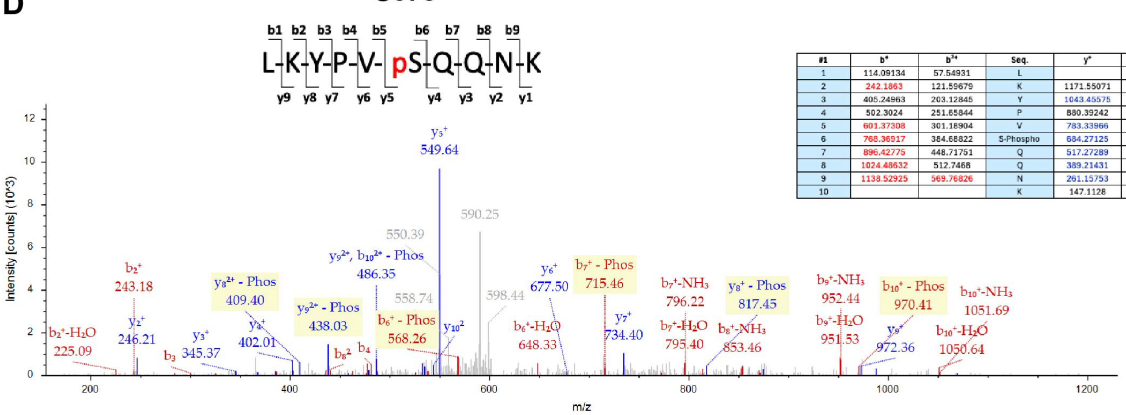


| #1 | b⁺ | b²⁺ | Seq. | y⁺ | y²⁺ | #2 |
|---|---|---|---|---|---|---|
| 1 | 114.09134 | 57.54931 | L | | | 10 |
| 2 | 242.1863 | 121.59679 | K | 1171.55071 | 586.279 | 9 |
| 3 | 405.24963 | 203.12845 | Y | 1043.45575 | 522.23151 | 8 |
| 4 | 502.3024 | 251.65844 | P | 880.39242 | 440.69985 | 7 |
| 5 | 601.37308 | 301.18904 | V | 783.33966 | 392.17347 | 6 |
| 6 | 768.36917 | 384.68822 | S-Phospho | 684.27125 | 342.63926 | 5 |
| 7 | 896.42775 | 448.71751 | Q | 517.27289 | 259.14008 | 4 |
| 8 | 1024.48632 | 512.7468 | Q | 389.21431 | 195.11079 | 3 |
| 9 | 1138.52925 | 569.76826 | N | 261.15753 | 131.0815 | 2 |
| 10 | | | K | 147.1128 | 74.06004 | 1 |

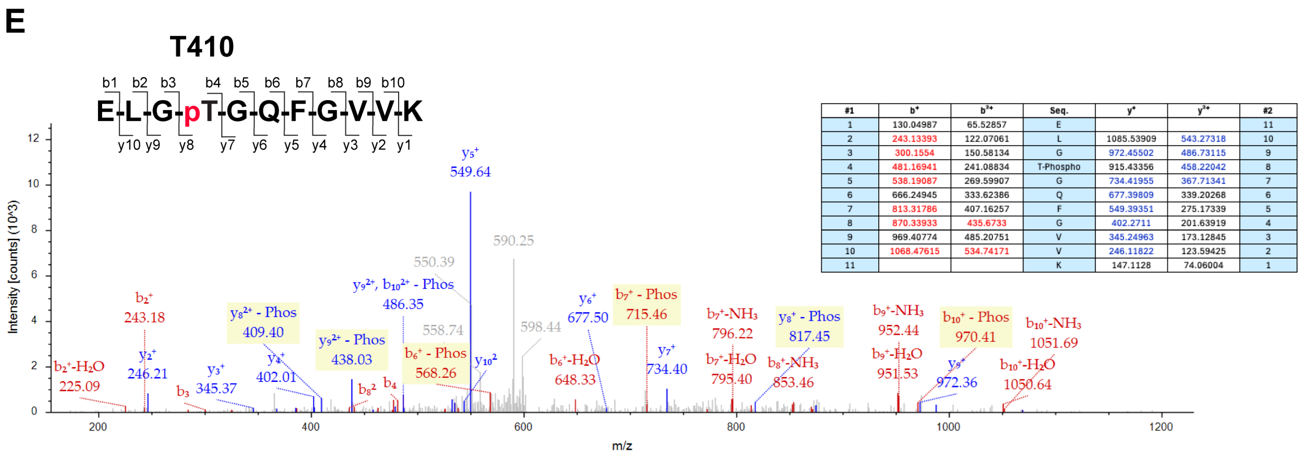


| #1 | b+ | b2+ | Seq. | y+ | y2+ | #2 |
|---|---|---|---|---|---|---|
| 1 | 130.04987 | 65.52857 | E | | | 11 |
| 2 | 243.13393 | 122.07061 | L | 1085.53909 | 543.27318 | 10 |
| 3 | 300.1554 | 150.58134 | G | 972.45502 | 486.73115 | 9 |
| 4 | 481.16941 | 241.08834 | T-Phospho | 915.43356 | 458.22042 | 8 |
| 5 | 538.19087 | 269.59907 | G | 734.41955 | 367.71341 | 7 |
| 6 | 666.24945 | 333.62386 | Q | 677.39809 | 339.20268 | 6 |
| 7 | 813.31786 | 407.16257 | F | 549.39351 | 275.17339 | 5 |
| 8 | 870.33933 | 435.6733 | G | 402.2711 | 201.63919 | 4 |
| 9 | 969.40774 | 485.20751 | V | 345.24963 | 173.12845 | 3 |
| 10 | 1068.47615 | 534.74171 | V | 246.11822 | 123.59425 | 2 |
| 11 | | | K | 147.1128 | 74.06004 | 1 |

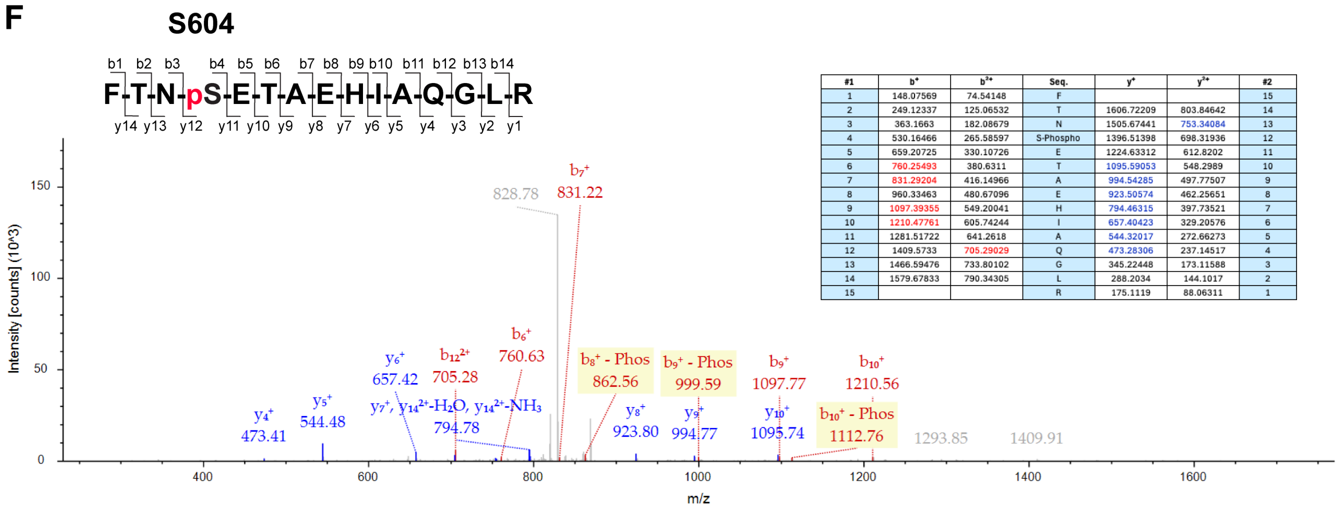


| #1 | b+ | b2+ | Seq. | y+ | y2+ | #2 |
|---|---|---|---|---|---|---|
| 1 | 148.07569 | 74.54148 | F | | | 15 |
| 2 | 249.12337 | 125.06532 | T | 1606.72209 | 803.84642 | 14 |
| 3 | 363.1663 | 182.08679 | N | 1505.67441 | 753.34084 | 13 |
| 4 | 530.16466 | 265.58597 | S-Phospho | 1396.51398 | 698.31936 | 12 |
| 5 | 659.20725 | 330.10726 | E | 1224.63312 | 612.8202 | 11 |
| 6 | 760.25493 | 380.6311 | T | 1095.59053 | 548.2989 | 10 |
| 7 | 831.29204 | 416.14966 | A | 994.54285 | 497.77507 | 9 |
| 8 | 960.33463 | 480.67096 | E | 923.50574 | 462.25651 | 8 |
| 9 | 1097.39355 | 549.20041 | H | 794.46315 | 397.73521 | 7 |
| 10 | 1210.47761 | 605.74244 | I | 657.40423 | 329.20576 | 6 |
| 11 | 1281.51722 | 641.2618 | A | 544.32017 | 272.66273 | 5 |
| 12 | 1409.5733 | 705.29029 | Q | 473.28306 | 237.14517 | 4 |
| 13 | 1466.59476 | 733.80102 | G | 345.22448 | 173.11588 | 3 |
| 14 | 1579.67833 | 790.34305 | L | 288.2034 | 144.1017 | 2 |
| 15 | | | R | 175.1119 | 88.06311 | 1 |

**Figure S6**. Mass spectrometry-based identification of BTK phosphorylation induced by AMPK-BTK PHICS. Spectra for the peptides with phosphorylated T191 (**A**), T316 (**B**), S366 (**C**), S378 (**D**), T410 (**E**), and S604 (**F**). The fragmentation pattern is shown in each spectrum.

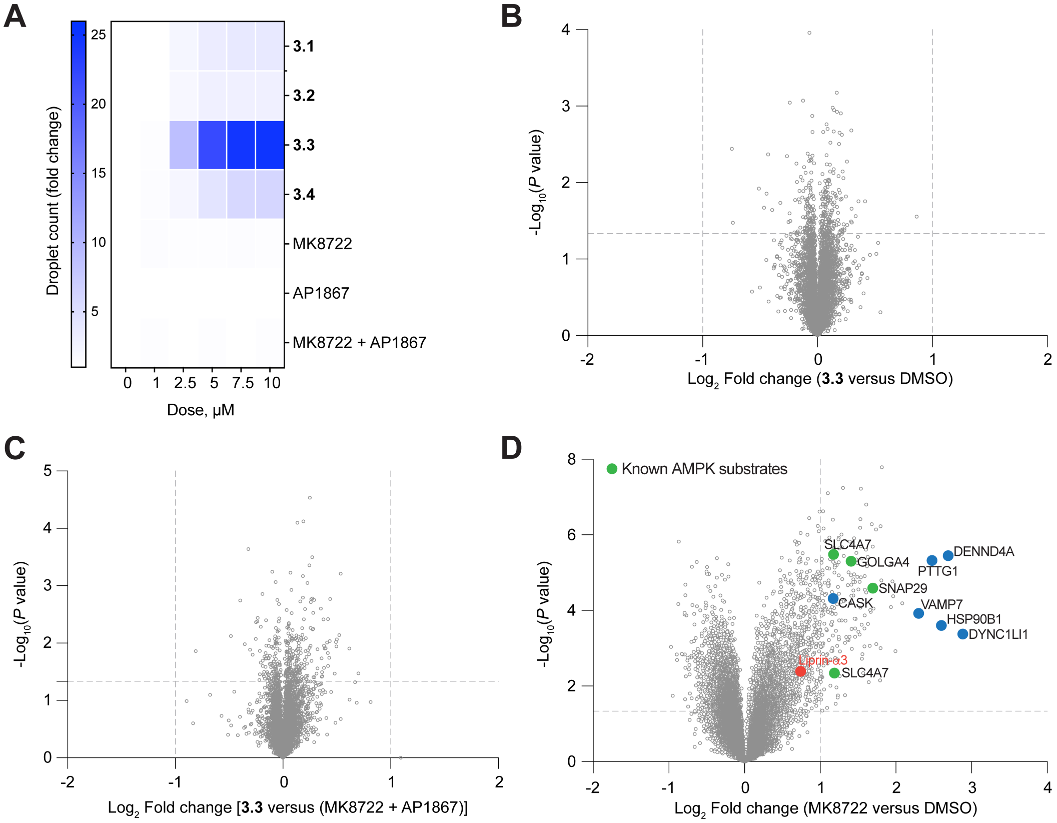


**Figure S7**. Liprin-α3 phase separation induced by AMPK-FKBP12$^{F36V}$-PHICS mediated phosphorylation. (**A**) Dose-dependent induction of Liprin-α3 condensates by PHICS and individual binders. (**B, C**) Global protein expression induced by **3.3** (5 μM) compared to DMSO (B), or equimolar (5 μM) combination of MK8722 and AP1867 (C). (**D**) Volcano plots displaying the $-\log_{10}$ (*P*-value) versus $\log_2$ fold-change of phosphopeptides by MK8722 versus DMSO.
